# LC3 forms functional nanocluster on autophagosome

**DOI:** 10.1101/2024.12.21.629869

**Authors:** Saman Fatihi, Deepak M. Khushalani, Joydipta Kar, Nitin Mohan, Lipi Thukral

## Abstract

Autophagosome biogenesis relies on the intricate coordination of proteins and lipids, with LC3B proteins persistently anchored to the double-membraned autophagosomes through a lipid anchor. However, little is known about how LC3B is organized in high concentration, its spatial distribution, and the mechanisms underlying its protein-mediated tethering on membranes. Using molecular dynamics simulations and super-resolution microscopy, we demonstrate that LC3B self-assembles to form higher-order clusters, averaging 150 nm in size. Interestingly, simulations provided cue for LC3B and phosphatidylinositol lipid specificity, and STORM imaging confirmed a clear overlap of LC3B-enriched regions with phosphatidylinositol-3-phosphate lipids. Together, LC3B nanoclusters on these lipids form spatially distinct “islands” on the autophagosome. Additionally, molecular analysis of 296 clusters revealed that clustering is driven by a unique rear binding pocket in Loop6 defined by alternating hydrophobic and polar residues. We generated four mutants to disrupt the characteristic self-assembly motif, with all four mutants resulting in aberrant cluster formation and impaired autophagosome motility. These findings highlight that LC3B self-assembly is crucial for autophagy and suggest a spatiotemporal mechanism regulating LC3B function.

## Introduction

The autophagosome is a unique double-membrane vesicle formed entirely *de novo* during the process of autophagy [1,2]. The biogenesis of autophagosomes is tightly regulated by a complex interplay of autophagy-related proteins, predominantly driven by their interactions with membrane lipids [3,4]. Central to this process is the LC3B protein, which plays a crucial role at multiple stages of autophagy [5]. Upon autophagic induction, LC3B translocate to the growing phagophore, where it facilitates phagophore elongation, autophagosome maturation, and ultimately facilitates fusion with lysosomes for cargo degradation [6,7]. A key step in this process is the conjugation of cytoplasmic LC3B to phosphatidylethanolamine (PE), producing the membrane-bound form LC3B-II [8–10]. The lipidated form of LC3B specifically associates with autophagosomal membranes, serving as a key marker of autophagic activity. Its presence in punctate, spot-like appearance reflects the abundance and spatial localization of autophagosomes within the cell (11). Fluorescent labelling distinguishes between the cytosolic and membrane-bound forms of LC3B, where a diffuse cytosolic pattern indicates low autophagic activity, while increased puncta number and size signify elevated autophagic flux [12]. While the interaction of LC3B with phosphatidylethanolamine (PE) and phosphatidylserine (PS) through lipid anchoring is well-established, the dynamics of LC3B and its interactions with other membrane lipids remain poorly understood [13]. Thus, there exists significant gap in our understanding of how LC3B proteins are organized on diverse array of lipids on the autophagosome membrane. This gap arises partly from the challenges inherent in studying small membrane-associated proteins using structural biology techniques, as well as the lack of a molecular model that elucidates the organization of concentrated lipidated LC3B proteins on membranes. To bridge this gap, we employed a combination of molecular simulations and super-resolution imaging to explore the dynamic organization of LC3B tethered to membranes via a lipid anchor under crowded conditions.

Recent advances in super-resolution microscopy have surpassed the diffraction limit of conventional light microscopy, enabling visualization of cellular structures and molecular interactions at the nanometer scale [14]. Single molecule localization techniques such as Stochastic Optical Reconstruction Microscopy (STORM) and Photo Activated Localisation Microscopy (PALM) provide lateral resolution of ∼20 nm, making them ideal for studying protein clusters [15], large molecular complexes [16,17], and detailed visualisation of cellular ultrastructure [18,19]. Further the nanometer resolution also allows precise quantification of stoichiometry, as well as relative positioning of membrane bound proteins on cellular organelles [20]. Recently, with Structured Illumination Microscopy (SIM), with ∼100 nm resolution, it was shown that autophagosome precursors emerge from recycling endosomes with complex morphologies, revealing finger-like structures that mature into spherical forms [21]. However, studying the dynamic organization of proteins at atomistic level on membranes remains a challenge. Molecular dynamics (MD) simulations have emerged as a powerful tool to complement these limitations, providing rich insights into biological interactions, particularly protein-lipid dynamics. Small peripheral membrane proteins, like LC3B, provide an excellent opportunity for investigation through these complementary approaches, offering insights into their dynamics, interactions, and functional roles with high precision.

The structural understanding of LC3 has significantly advanced in recent years, shedding light on its multifaceted roles in autophagy [22–24]. Its structure consists of an N-terminal helical region and a ubiquitin-like domain that provides sites for interaction with autophagy adaptors and membrane components. LC3 features two canonical hydrophobic binding pockets that enable LC3 Interacting Region (LIR)-mediated interactions with autophagy receptors, such as p62, NBR1, and optineurin, bridging LC3 with cargo destined for degradation. Human LC3 proteins consist of six isoforms: LC3A, LC3B, LC3C, GABARAP, GABARAPL1, and GATE16. They are closely related yet perform distinct functions [25]. Our previous work on LC3B (hereafter referred to as LC3) has elaborated on its membrane-targeting motif [26], its spontaneous binding process during lipidation [27], and the role of LC3 human homologs in shaping diverse functional outcomes [28].

In this study, we combined coarse-grain molecular simulations with nanoscopic super-resolution imaging to extract two layers of information. First, we investigated the organization of LC3 at higher concentrations on membrane lipids. Through multiple independent simulations, we identified the formation of higher-order clusters of LC3 and a remarkable preponderance of LC3 to be associated with phosphatidylinositol (PI) lipids on a heterogenous lipid membrane. This observation was consistent with STORM imaging, where nanometer-sized LC3-rich regions were detected across the autophagosomes. Interestingly, these nanoclusters were organized on phosphatidylinositol-3-phosphate (PI3P) lipids, forming distinct spatially uncoupled ‘LC3 islands’ on the autophagosomes. Secondly, we sought to understand the molecular mechanisms driving LC3 clustering, i.e., identifying which forces or regions were involved and whether the protein clusters are driven by specific or non-specific interactions. By comparing wild-type LC3 with four engineered mutants designed to disrupt cluster formation, we identified a novel rear binding surface on the LC3 protein. The region was characterized by alternative polar and non-polar amino acids, commonly associated with self-assembly of proteins. These molecular insights, combined with functional assays of autophagosome motility, suggest that LC3 clusters, in conjunction with PI lipids, are involved in spatiotemporal regulation in autophagy.

## Results

### Coarse-grain molecular simulations show clusters of lipidated LC3 on autophagosomal membranes

To understand LC3 organisation on membrane in a crowded environment, we generated a physiologically relevant model with multiple lipidated LC3 proteins on heterogenous membrane mimicking autophagosomal lipids. Figure 1A shows the simulation system, consisting of 90 LC3 proteins inserted via PE chain into a lipid bilayer of 100 nm length. All molecules are represented at a coarse-grained representation, with approximately 1 million atoms. The bilayer is composed of mixture of phosphatidylcholine (PC), phosphatidylethanolamine (PE), phosphatidylserine (PS) and phosphatidylinositol (PI) lipids reflecting the ER composition that largely mimics the lipid source of the autophagosome. The LC3 proteins were uniformly distributed, with each protein spaced roughly 8 nm apart, covering 26% of the membrane surface. We observed that LC3 proteins formed higher-order clusters by the end of the initial simulation (Figure 1B). To validate this observation, two additional simulations, each 10 μs long, were performed. Across all replicas, a consistent tendency for all proteins to cluster was observed (Figure 1C, S1). Dimers initially formed within 500 ns, and by the end of the simulation, nearly all LC3 molecules were clustered (Figure S2). The interactions were predominantly limited to two neighbouring LC3 proteins, although extensive interfaces involving three or more partners were also observed (Figure 1D, S3). All LC3 clusters were properly conjugated to the membrane, with the PE anchor consistently embedded within the headgroups of the upper leaflet for each LC3 protein throughout the simulation (Figure 1E).

**Figure 1.**
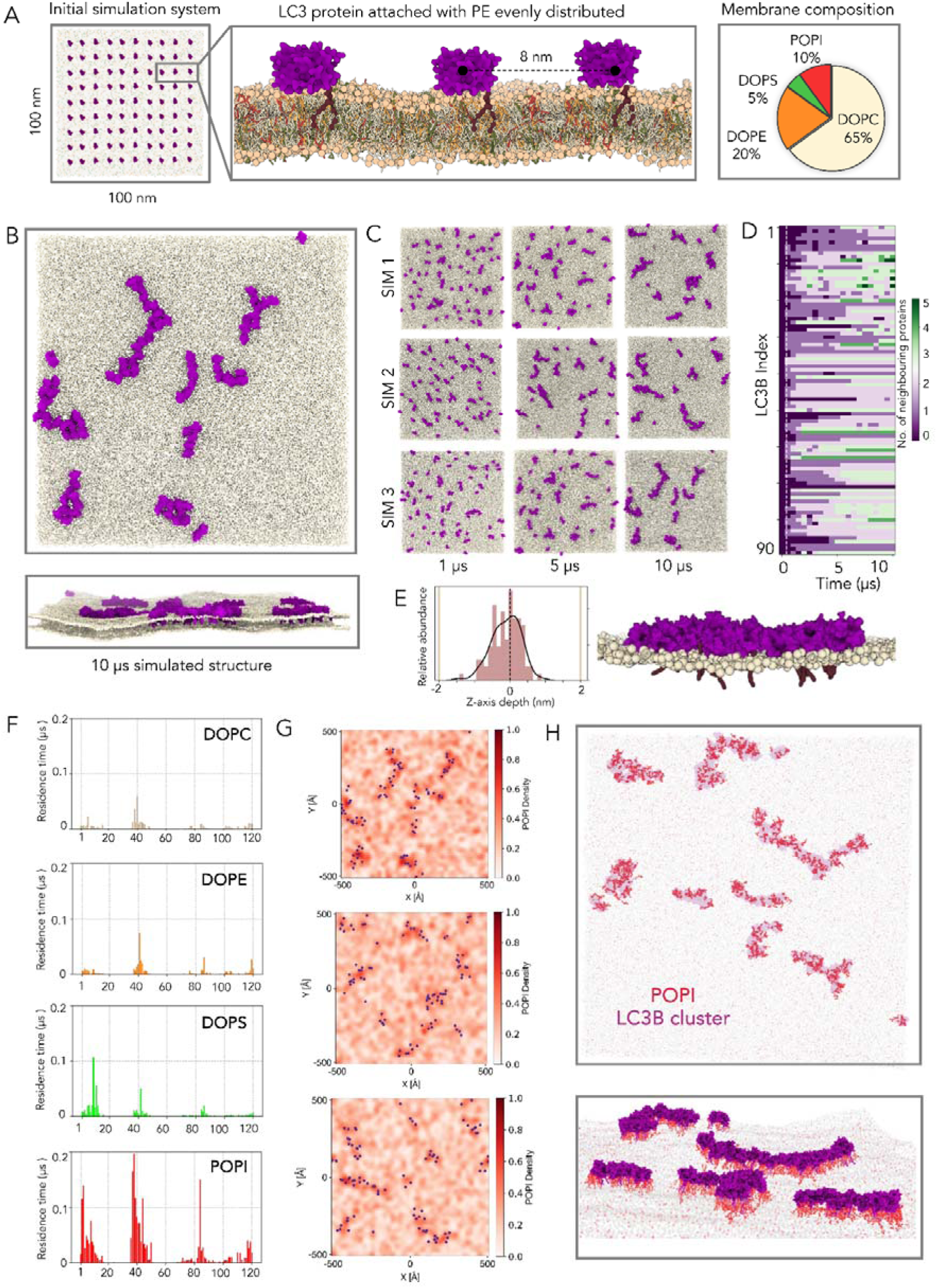
Lipidated LC3 self-assemble to forms clusters on autophagosomes. A) The snapshot shows the initial setup with 90 LC3B molecules evenly distributed on the outer leaflet of the membrane. The protein is covalently attached with PE and are evenly distributed at a distance of 8 nm apart. The membrane is composed of heterogenous ER lipids, including DOPC, DOPE, DOPS, and POPI. B) Representative snapshots shown depicting the formation of non-uniform LC3B nanoclusters in top and front view. C) All the replicas showed clear tendency to self-assemble, as seen in three time points for three coarse-grain simulations. D) Number of neighbouring LC3 proteins in a given cluster is calculated across all 90 proteins. E) The density plot shows the distribution of PE anchor insertion for each LC3B molecule in the outer leaflet headgroups. The zoomed in snapshot highlights the orientation of representative clusters on the membrane with PE-tail inserted inside the PO4 headgroups showcasing the clustering of proteins observed on the membrane. F) Residence time for each lipid is calculated in proximity of residues in LC3 proteins. G) Probability distribution of lipid density calculated across simulation XY plane. The dots represent the LC3 proteins, a significant overlay is observed with PI lipids. H) The front and top view of accumulation of PI lipids in proximity to LC3 clusters.

The key feature of membrane targeting module of LC3 is highly enriched with eleven positively charged residues, as also shown in our previous work [26]. We found consistent pattern of these residues, especially from the N-terminal and loop regions, suggesting electrostatic interactions with the autophagosomal membrane (Figure S4). To test this hypothesis, we computed lipid interactions in proximity to LC3 clusters. While phosphatidylcholine (PC) and phosphatidylethanolamine (PE) are the most abundant lipids, comprising approximately 65% and 20% of the total phospholipids, respectively, LC3 interacts strongly with low-abundant PI lipids. We calculated the number of contacts between different lipids and each residue of the protein, normalizing these values by lipid concentration to enable comparative analysis (Figure 1F). Amongst the charged residues within membrane targeting module, the inositol groups showed specific interactions with three positively charged residues (Arg37, Lys39, and Lys42) located at the start of loop 3 of LC3 (Figure S5). Probability density plots from three simulations revealed a significant enrichment of PI lipids around the LC3 clusters compared to other lipids (Figure 1G, Figure S6). Closer examination of these clusters showed clear traces of PI lipids concentrated at the centre and in proximity to the clusters, far exceeding the levels of other phospholipid moieties (Figure 1H). Our findings show that LC3 conjugated with PE has strong tendency to self-assemble on membrane and form higher-order oligomers that show high enrichment with PI lipids.

### Super-resolution microscopy confirm formation of LC3 nanoclusters of autophagosomes

Observations from the simulations prompted us to experimentally resolve and validate the self-organization of LC3 on autophagosomes. Towards this, we performed super-resolution STORM imaging of LC3 in starved BS-C-1 cells (Figure 2A-B). While conventional TIRF imaging of LC3 depicts diffraction-limited autophagosome puncta within the cell (Figure 2A inset; Figure S7), 3D STORM imaging allows us to resolve the spatial distribution of LC3 on each autophagosome. At 20 nm spatial resolution, we visualize that membrane-bound LC3 forms spatially distinct clusters, spanning the autophagosomal membrane (Figure 2C). Quantitative analysis of LC3 clusters using DBSCAN, enumerates significant heterogeneity in the number of clusters, with autophagosomes containing 5-25 LC3 clusters, and an average of 9 LC3 clusters per autophagosome (Figure 2D). We hypothesized that the heterogeneity in LC3 cluster numbers is driven by the variability in autophagosome size, which ranges from 500 nm to 3 μm in diameter within cells (Figure S8). To test this, we analyzed the relationship between LC3 cluster numbers and autophagosome size, observing a clear linear correlation (Figure 2E). This confirms that larger autophagosomes can accommodate a greater number of LC3 clusters.

**Figure 2:**
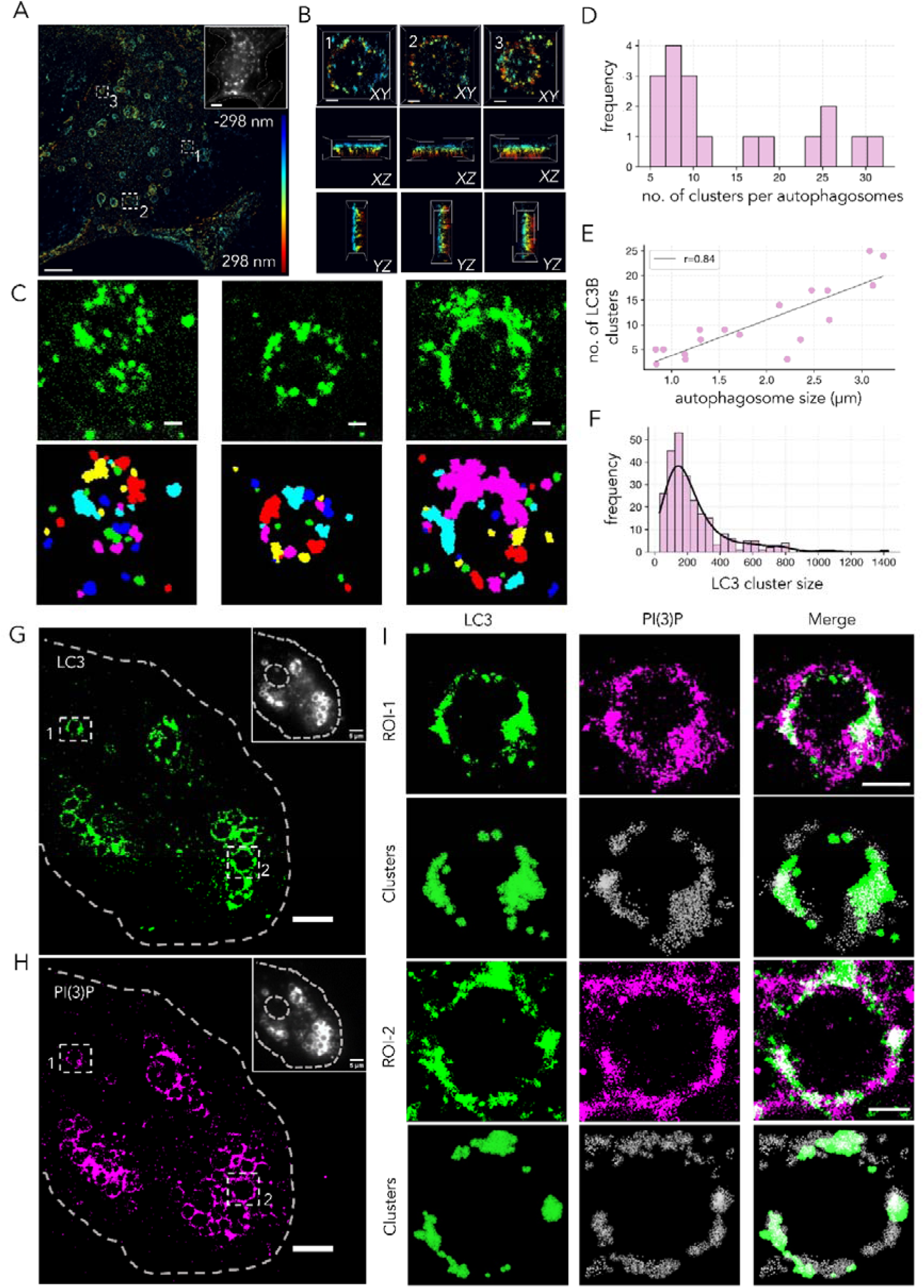
TIRF and STORM imaging of autophagosomes showing LC3 nanoclusters. A) Super-resolution 3D STORM imaging of starved BS-C-1 cells, immunostained for endogenous LC3B. Inset is the diffraction-limited TIRF image of the same cell highlighting LC3B puncta formation on autophagosomes. B) Three different regions of interest (ROIs) from image A, demonstrating the spatial organization of LC3B on mature autophagosomes in XY, XZ, and YZ planes, highlighting LC3B clusters spanning the membrane. C) 2D STORM images of LC3B decorated on autophagosomes of varying sizes across three different ROIs, along with corresponding DBSCAN-segmented images, reveal the highly resolved spatial distribution of LC3B organized into nanoclusters on autophagosomes. In the segmented image colors represent distinct nanoclusters. D) Quantitative cluster analysis of LC3B on autophagosomes is shown with number of LC3B nanoclusters on each autophagosome. E) Correlation between the number of clusters and autophagosome size. F) Size distribution of LC3B nanoclusters. G-H) Sequential 2-colour 2D STORM imaging of starved BS-C-1 cells overexpressing the 2X FYVE domain of HRS protein tagged with mCherry immunostained for endogenous LC3B, and for mCherry. Respective insets are the diffraction-limited TIRF image of the LC3 (inset of G)-and mCherry (inset of H) in the same cell. I) Two different regions of interest (ROIs) from image G and H with corresponding DBSCAN segmentation analysis, demonstrating the spatial organization of LC3B on autophagosomes enriched with PI(3)P. The LC3B cluster organization exhibits remarkable similarity to the PI(3)P nano-domain architecture on the autophagosomes. Scale bars: (C) and (I) 200 nm; (G) and (H) 5 µm.

Next, when we analyse the size of the LC3 clusters, we observed a peak size distribution of LC3 clusters at 150 nm. Albeit, about 10% of the population exhibit clusters of extremely large cluster size (>400 nm) contributed from the 10% of heterogeneity in the cluster size on each autophagosome (Figure 2F). Interestingly, unlike cluster number, LC3 cluster size does not scale linearly with autophagosome size (Figure S9). Instead, LC3 clusters maintain a consistent size range of 100–300 nm, regardless of autophagosome dimensions. To adapt to the available autophagosomal membrane surface, LC3 clusters do not grow larger but instead increase in number, preserving their optimal size range for effective autophagosome decoration.

Among all phosphoinositides, PI(3)P is well-established as a key player in autophagy initiation [29–31]. Therefore, we transiently overexpressed a PI(3)P lipid sensor, mCherry tagged tandem-FYVE domain of the HRS protein, which specifically binds to membranes decorated with PI(3)P lipids. Cells were starved for 2.5 hours, followed by a sequential immunostaining and 2D STORM imaging, as detailed in the methods. Strikingly, we observed that PI(3)P lipids also form spatially distinct nanodomains on the autophagosome surface (Figure 2G-H). Furthermore, LC3 clusters were found to colocalize with PI(3)P nanodomains. Image segmentation using DBSCAN revealed a remarkable similarity in the shape and pattern that represents spatial organization and localization distribution pattern of LC3 clusters and PI(3)P nanodomains (Figure 2I). These findings are consistent with our simulation results, indicating that the formation of LC3 oligomers is closely associated with the presence of specific signalling lipids, such as PI, on autophagosomes. Collectively, this demonstrates that LC3 organizes into dense, and spatially distinct nanoclusters enriched with specific PI3P lipids on the autophagosomal membrane.

### Identification of residues in the clustering binding interface

The simulations provided a unique opportunity to investigate interface properties across a diverse array of LC3 protein clusters engaged with the membrane. Across three independent 10 µs simulations, approximately 30 clusters were extracted from the final trajectories, yielding a comprehensive dataset of 296 distinct interacting surfaces (Figure S10). The analysis revealed that 95% of contact points originated from α1, α2, loop6, α4, and the C-terminal regions, forming two structurally contiguous motifs despite being dispersed in sequence space (Figure 3A, Figure S11). The first interface originates from loop6 in the core region, with peripheral residues from α4 and the C-terminal regions surrounding it. The second interface is in the N-terminal regions of both helices (α1-2) and loop3, which is partially membrane-embedded. For peripheral membrane proteins like LC3, interactions are often transient, contributing to heterogeneity in their binding modes. From above two interfaces, four heterogeneous dimer combinations were identified (Figure 3B). Structural analysis revealed that the N-terminal front region of these interfaces engaged in short-lived, transient interactions. In contrast, time-evolution data for four interaction pairs showed that residues in Loop6 consistently formed stable contacts with all clusters within 200 ns, acting as an early interaction hub (Figure S12).

**Figure 3.**
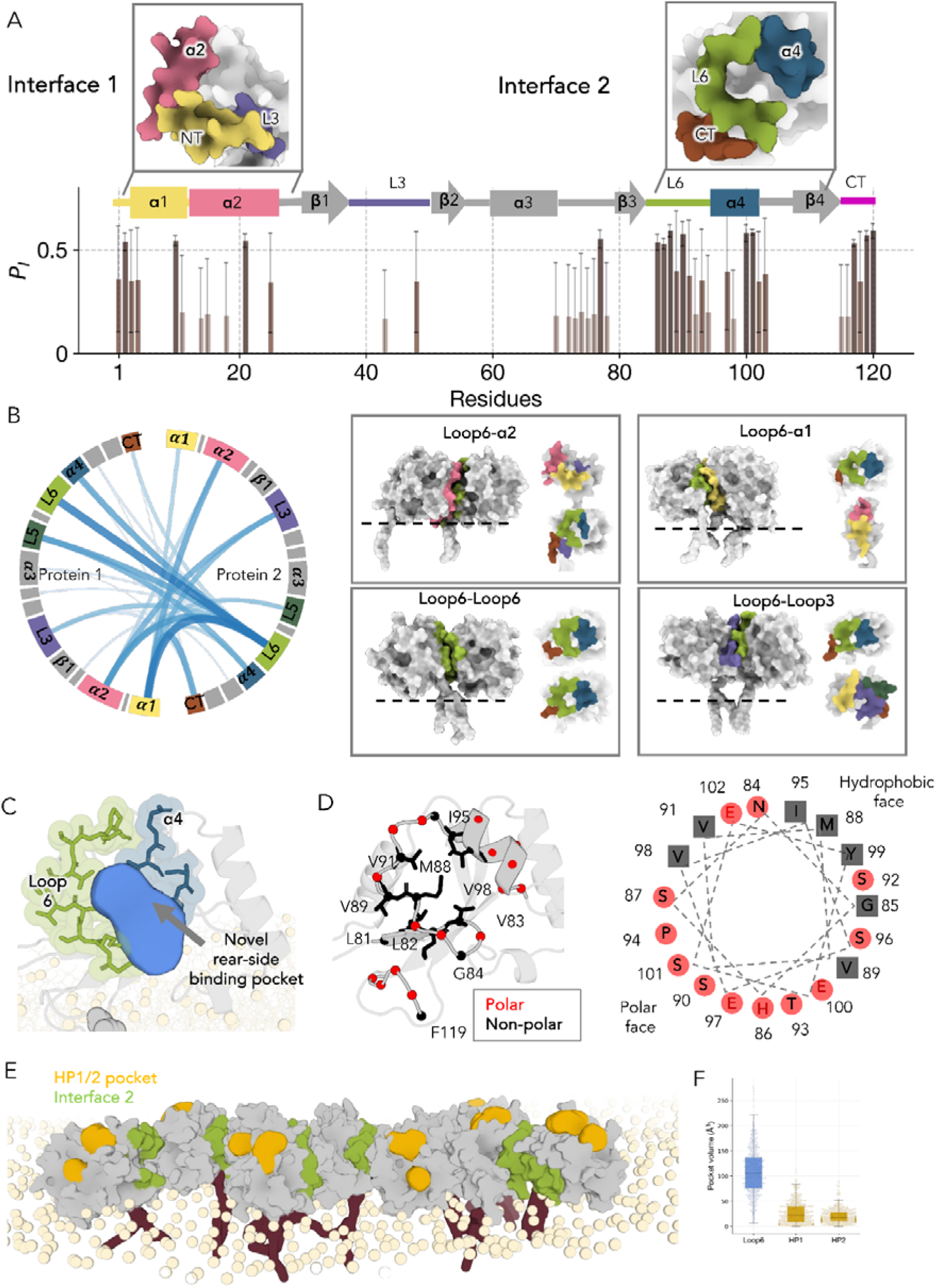
Identification of membrane-bound LC3-PE clustering interface. A) Residue-wise interaction probability plot depicting the likelihood of each residue to be at the interface across all three simulations. The contact is defined with a distance cutoff at 10 Å and >30% duration during the length of the trajectory. Snapshots of the two most probable contiguous interfaces (Interface 1 and 2) are depicted in surface representation. B) Circos plot showing interaction between 296 dimer interfaces, with width of edges directly proportional to number of contacts. From the dimer-dimer interface calculation of interface 1 and interface 2, four heterogenous dimer combinations are identified shown in boxes. C) The rear side of the protein was involved in most probably interface which also forms a significant cavity/pocket shown in blue colour. D) The novel binding cavity had a surface fingerprint that showed alternating polar and hydrophobic residues, a pattern found in many self-assembly proteins. Alongside, individual residues are shown for clarity of the non-polar and polar wheel. E) One of the higher-order cluster with concomitant marking of canonical hydrophobic receptor binding pockets and clustering interface at rear side that are non-overlapping. F) The bar plot showing surface accessibility of three pockets.

We then asked a question what molecular features make these contacts persistently sticky? As shown in Figure 3C, structural analysis revealed that interface 2 residues form a distinct pocket, with loop6 and α4 contributing to the formation of a “rear binding pocket”. There are three notable features of this pocket. First, in 43 experimentally solved LC3 complexes with other peptides or proteins, this region is not occupied, suggesting a potential interface for self-interaction (Figure S13). Second, an analysis of the properties of this region revealed an intriguing pattern of alternating polar (S87, S90, S91 ,T93) and hydrophobic residues (M88, V89, V91) forming a sticky interaction module (Figure 3D). Interestingly, this pattern aligns with the “binary code” concept previously identified in protein-protein interaction interfaces, where alternating polar and non-polar residues facilitate stable interactions by promoting both specific binding and adaptability at the interface, supporting functional protein interactions [32–34]. Also, the residues constituting this pattern are conserved, including “SMVSVS” (Figure S14). We hypothesized that for a small protein to form clusters, residues would need to be highly accessible, allowing adjacent pairs to assemble more easily. To perform this calculation, we backmapped coarse-grained simulations to atomistic resolution and minimized the structures for 100 ns (Figure S15). We used structures both prior to clustering, where no inter-protein binding occurred, and post-clustering, to calculate the surface accessibility of interfacial residues. Nearly 50% of residues in clustering interfaces were accessible. Lastly, this rear-side binding pocket is distinct from the canonical hydrophobic HP1 and HP2 binding pockets (Figure 3E). Across simulations, all residues involved in receptor interactions remain surface-accessible, confirming their potential for receptor engagement (Figure S16). In contrast, the rear binding pocket, with a volume of 115 Å³, facilitates the ability of LC3 proteins to self-assemble on the membrane (Figure 3F). Our findings identify a novel binding surface on the rear side of LC3, composed of alternating polar and hydrophobic residues, which drives the clustering of LC3 at membrane interfaces.

### Generating LC3 mutants from simulations to alter clustering interface

To understand the specific contributions of polarity and hydrophobic interactions driving LC3 cluster formation, we systematically introduced mutations to alter the clustering binding pocket. A total of four mutants were generated (Figure 4A, Figure S18). In mutant 1, we increased the polarity of the Loop6 region by substituting the hydrophobic residues with serine, namely three substitutions were made (M88S, V89S, and V91S). The second mutant 2 we focused on serine and altered the hydrophobicity within Loop6 (S87A, S90A, S92A, T93A, and P94A). Furthermore, we designed two negative control mutants in C-terminal and loop3 that may interfere non-specifically to disrupt regions peripheral to the specific clustering surface, ensuring minimal disturbance to the clustering interface. To understand their dynamics, we generated four mutant dimer simulations in atomistic detail to fully capture sidechain dynamics hitherto not fully understood in coarse-grained resolution. Simulations of mutant variants will also provided valuable insights into the role of specific residues in maintaining the structural integrity and dynamics of LC3 clusters. In our simulations, the wild-type dimer maintained a stable clustered conformation throughout, while both Loop6 (L6) mutations exhibited clear dissociation of clustering behavior (Figure 4B-C), indicating disruption in structural stability. Specifically, mutant 1 showed instability, dissociating within 100 ns, whereas mutant 2 dissociated at 400 ns. In contrast, both interface 1 derived mutants maintained their cluster formation (Figure S19, S20). These findings highlight the role of forces driving cluster formation, as accurately captured in physics-based simulation models.

**Figure 4.**
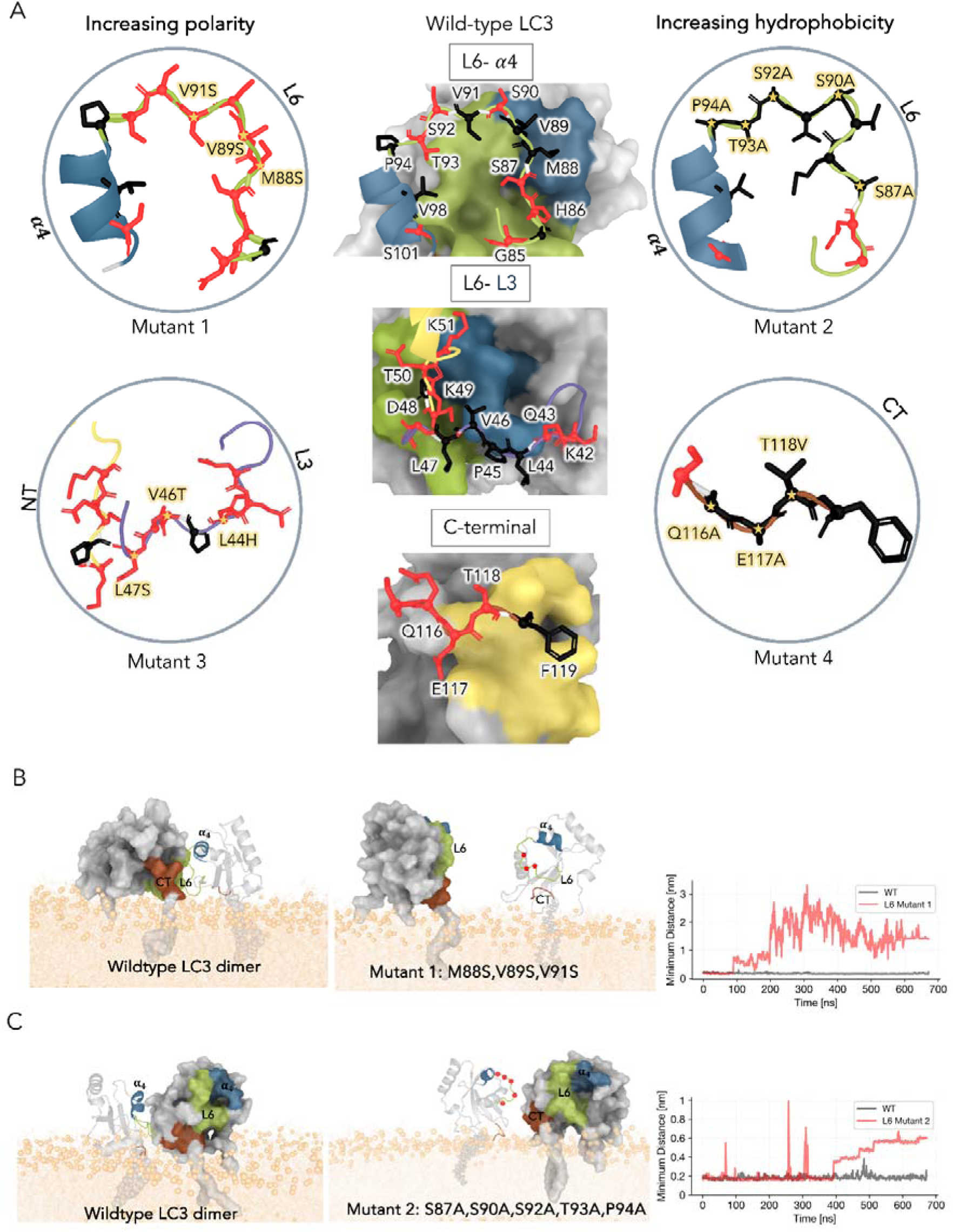
Designing mutants to abrogate LC3B clustering. A) Structural representations of the wild-type LC3 interfaces are highlighted. Key residues at the interface were targeted for mutagenesis to disrupt clustering. Zoomed-in structural models display the side chains of the designed mutants, illustrating changes in properties, either increased polarity (Loop6_M1: M88S, V89S, V91S; Loop3: L44H, V46T, L47S) or increased hydrophobicity (Loop6_M2: S87A, S90A, S92A, T93A, P94A; C-terminal: Q116A, E117A, T118V), with the mutated residues marked by star and labeled. B-C) Structure snapshots embedded membrane for two mutants (mutant 1 and mutant 2) after 650 ns of molecular dynamics simulations, alongside the wild-type (WT) in the same starting configuration is shown. The mutant exhibit increased separation between LC3 molecules, indicating clustering disruption. The right panel displays the time evolution of the minimum inter-molecular distance between the two LC3B molecules.

### STORM imaging of LC3 mutants shows altered clustering pattern and impaired motility

From above molecular dynamics simulations, we obtained dynamic consequences of mutations introduced in the clustering interface. To experimentally validate these mutations and understand their clustering behaviour on autophagosome, four mutant constructs were generated. Each LC3 mutant variant was tagged with eGFP at the N-terminal to distinguish the mutant populations from the endogenous population when expressed in the cells. To specifically understand the impact of mutations on the formation of nanoclusters, we transiently overexpressed LC3 wild-type and mutant variants tagged with eGFP in BS-C-1 cells, starved them, immunostained for eGFP and performed STORM imaging followed by DBSCAN based cluster analysis (Figure 5A). Our positive control is wild-type LC3-EGFP that showed similar cluster parameters as that of the endogenous population, while the two Loop 6 mutants shows no proper clustering, where LC3 appears to be evenly distributed on the autophagosome (Figure 5B and C). Hence, Loop 6 is one of the crucial regions for regulating LC3 cluster property. Interestingly, the typical spatially uncoupled LC3 cluster pattern observed in wild-type LC3 are notably distinct in the LC3 loop6 variants. We quantified the percentage of autophagosome area occupied by LC3, to observe that unlike the wild-type LC3, which forms nanoclusters concentrated on the autophagosome with 42% occupancy, the loop6 mutants exhibit a single, blob-like assembly on the membrane, covering approximately 80% of the autophagosomal surface. Furthermore, the two negative controls showed relatively less alteration in the clustering pattern with a shift in occupancy. The area of autophagosome occupied by LC3 clusters decreased by half in L3 & C-terminal mutant compared to the wild-type (Figure 5C), suggesting that the clusters formed are of smaller size. Also, multivalent interactions are often formed with lipids, which can make validation challenging. This data suggests that while these regions participate at the interface, they are not solely responsible for clustering.

**Figure 5.**
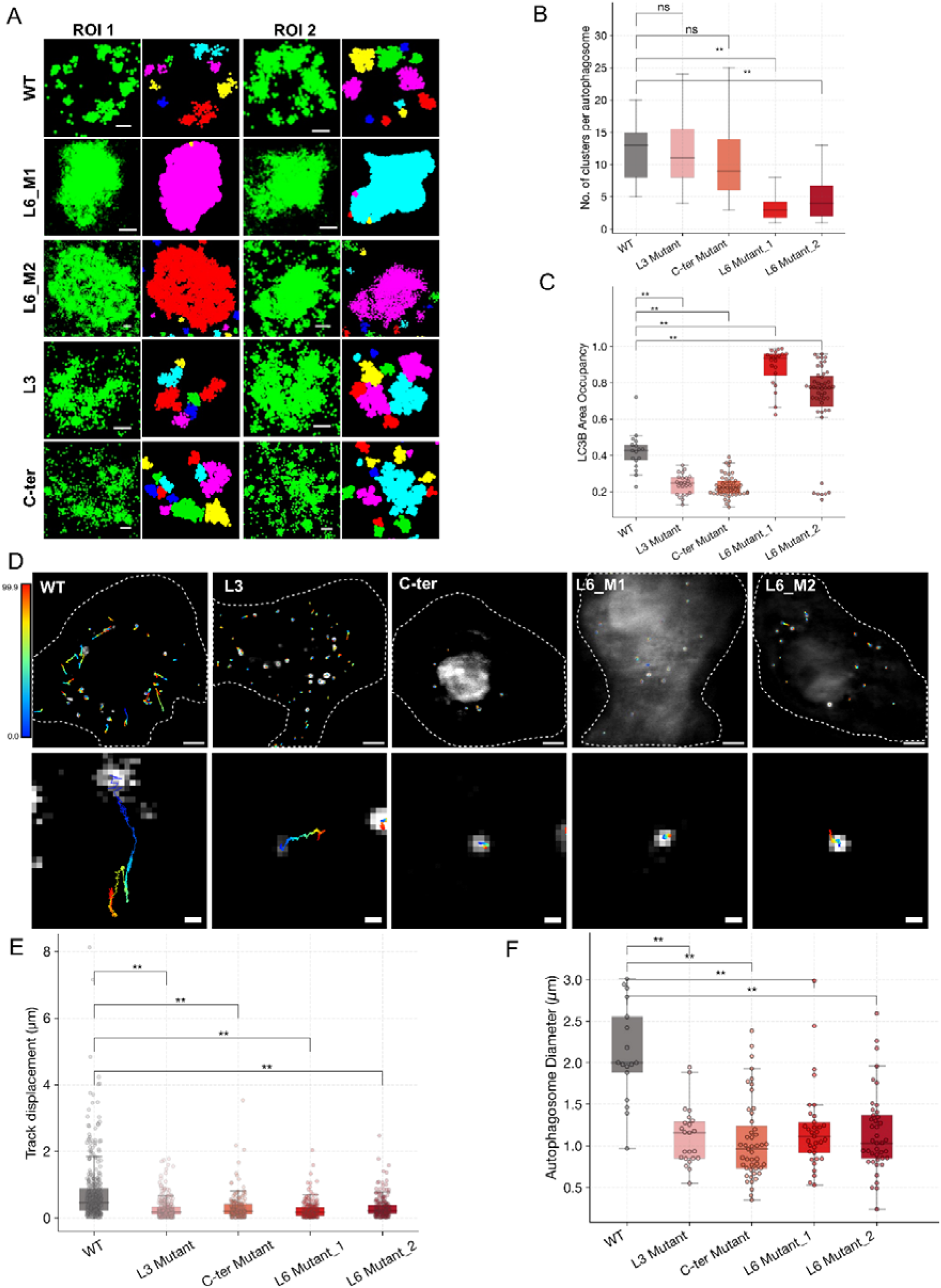
Super-resolution STORM images and cluster analysis of LC3B wildtype and mutants. A) Schematic showing the core and periphery region of the interface to have differential impact on LC3B clustering property. A) STORM images of single autophagosomes from cells transiently overexpressing LC3B_WT and indicated mutants in starved BS-C-1 cells with corresponding DBSCAN segmentation analysis. Segmented images are pseudo-colour-coded with different colours corresponding to different nanoclusters. Scale Bars 200 nm. B) number of LC3B clusters per autophagosome. C) Box plots showing the fraction of area occupied by LC3B on autophagosomes and D) Representative single particle tracking trajectories (colored lines) depicting the track length of EGFP-LC3B-positive autophagosome vesicles in BS-C-1 cells (20 fps). (Lower panel) Zoomed in shows magnified trajectory of a single autophagosome. The trajectories are color coded with respect to time scale (T). Scale bars 5 μm, zoom 500 nm. E) Distribution of track displacement for the wildtype and mutant LC3. F) Distribution of autophagosome size for wildtype and mutant LC3. All values indicated are mean of the respective distribution.

Newly formed and matured autophagosomes, guided by motor proteins, travel along microtubules to eventually fuse with lysosomes, forming autolysosomes [29,30]. Several studies have demonstrated that LC3 regulates the adaptor proteins necessary for recruiting and activating specific motors on autophagosomes, thereby controlling the unidirectional retrograde transport of autophagosomes within the cells [35,36]. To determine if LC3 oligomerization on the autophagosome membrane is essential for LC3 function, we investigated the motility of autophagosomes with varying levels of LC3 clustering induced by LC3 mutants. To this end, we performed live-cell imaging of BS-C-1 cells transfected with wild-type and mutant eGFP-LC3 variants subjected to starvation using TIRF microscopy (Figure 5D). Single-particle trajectory analysis of individual autophagosomes from live-cell imaging reveal that LC3 mutations lead to confined motion of the autophagosome, unlike the long-range directed transport observed with wild-type LC3 (Figure 5D lower panel). The track displacement analysis shows that autophagosomes decorated with wild-type LC3 exhibited long-distance movement in a typical directed manner, whereas those with mutant LC3 displayed hindered motility as seen by decrease in net displacement (Figure 5E). Since autophagosome motility was compromised for the Loop-6 mutant, which does not form LC3 clusters, as well as for the C-terminal and Loop 3 mutant, which forms smaller LC3 clusters, we conclude that an optimal LC3 cluster size and distribution are necessary for functional LC3 clustering. Interestingly, we also observe a remarkable decrease in autophagosome size with the mutant LC3 expression (Figure 5F). These results suggest a functional role of LC3 clustering in autophagosome formation.

## Discussion

The dynamic assembly of autophagosomes relies on reversible and transient interactions involving lipid-anchored LC3, densely packed on these double-membraned vesicles. The present understanding of LC3 has transitioned from a *molecular switch* oscillating between cytosolic LC3-I and membrane-associated LC3-II to being recognized as a key *facilitator* in the autophagic process. However, its presence on vesicular environments, characterized by their crowded nature, pose significant challenges for studying confined protein-protein interactions on highly heterogeneous membrane spaces. Also, high volume occupied media often leads to non-specific interactions and molecular noise, making it difficult to isolate biologically relevant interactions [37]. Previous report has indicated that LC3/Atg8 forms high molecular weight complexes upon conjugation with phosphatidylethanolamine (PE) [6], and prompted us to look into molecular self-assembly in greater detail.

In this work, we show that LC3 forms functional nanoclusters on autophagosome membranes packing optimally in spatially isolated islands. Using combined computational and experimental approach, we demonstrate that LC3 proteins assemble into nanometer-sized clusters optimally distributed on autophagosomes. Simulations also provided us the cue that there exists specific PI lipid enrichment on these clusters. STORM imaging of PI3P revealed a clear overlap with LC3 clusters, confirming the association between them. Molecular analysis revealed a novel rear binding pocket on LC3 proteins, characterized by an alternating polar and hydrophobic residues, which drives clustering. To disrupt this pattern, we generated four mutants: the first two targeting the Loop6 region, and the other two designed to minimally interfere with clustering. Extensive experimental and in silico mutant characterisation showed that disrupting the polarity and hydrophobicity of this pocket destabilizes the clusters, reducing their stickiness and impairing their stability.

By elucidating the molecular determinants of LC3 oligomerization on the membrane, our findings shed light on role of LC3 in the spatiotemporal regulation of autophagy. Several biological scenarios can be speculated, where LC3-enriched islands may serve as critical signalling platforms. The most plausible scenario is where LC3 clusters might be involved in the regulation of membrane dynamics, controlling the size and shape of autophagosomes. STORM imaging of all four mutants revealed smaller autophagosomes with disrupted cluster formation. While mutants 1 and 2 within Loop6 drastically altered the appearance of LC3 proteins, causing them to diffuse throughout the vesicle, the other two mutants showed only marginal changes in cluster size. It is possible that the heterogeneous clustering interface observed in our loop 6 interface 2, even when present in low abundance, is sensitive to control the size of the autophagosome. Findings from different experiments also show that LC3 plays an essential role in membrane dynamics [7,38], interacting with other autophagy-related proteins to aid in the elongation and closure of the phagophore [39].

Nishimura and Tooze review the emerging roles of various membrane lipids, such as phosphatidylinositol 3-phosphate (PI3P) and phosphatidylethanolamine (PE), and their interaction with proteins in autophagosome formation [40]. There are multiple autophagic proteins that have lipid-related roles. For instance, the distinctive shape of ATG17, as demonstrated structurally by Ragusa et al., resembles BAR proteins that induce membrane curvature, marking it as a unique membrane-lipid curvature sensing protein on autophagosomes [41]. Another lipid-related role in autophagy is from ATG9 vesicles and ATG2 proteins that act as essential lipid carriers in autophagy, providing key lipids for membrane formation and expansion [42]. Additionally, WIPI proteins and Atg13 have been shown to directly bind negatively charged lipids [43]. In particular, WIPI proteins are crucial for lipid recognition through their PI3P-binding FYVE motif [44]. These findings, along with our study showing the overlap of LC3 and PI lipids, further emphasize that the distribution of autophagy-related proteins could act as signalling platforms on unique lipids for the progression of autophagosome formation, ensuring that autophagy is precisely coordinated with cellular cues.

Also, key studies have highlighted how nanometer-scale assemblies of peripheral membrane proteins like lipid anchored small GTPase Ras proteins serve as platforms for membrane curvature, vesicle formation, and trafficking [45]. Such membrane bound proteins form transient signalling platforms that self-organize, often driven by lipid-lipid interactions [46], protein-lipid affinity [47], or external stimuli [48]. However, LC3 nanoclusters may also spatially and temporally control cargo degradation, and extend the role of peripheral proteins that are residing on plasma membranes. For instance, in the selective autophagy of specific organelles or protein aggregates, it may guide the autophagic machinery to the exact location where it is needed. By concentrating LC3 and other autophagy-related proteins in specific regions, these islands could help in the efficient recruitment of cargo and the orchestration of complex autophagic processes.

Our work also raises several important future questions about the nanoclusters themselves. How is the optimal spatial distribution of LC3 affected by changing environments, such as mechanical fusion with lysosomes or extracellular chemical stimuli? Additionally, how dynamics or movement of these nanodomains within lipid environments is controlled. Future studies by our groups will also explore understanding LC3 isoforms clustering, if any, and their association with autophagosome apparatus. Interactions of these nanoclusters with both autophagic and non-autophagic partners will provide insights into broader functional signalling networks and potential non-canonical roles of LC3 in cellular processes. The LC3 stoichiometry is also not fully aligned in this work with two approaches, as atomistic resolution reveals dimer and trimer populations forming before higher-order oligomers. However, the precise number of these molecules and their subsequent population distribution remains unknown. Further functional analysis will provide deeper insights into the regulatory mechanisms controlling these complexes and enhance our understanding of how LC3 oligomers contributes to autophagy.

## Materials and methods

### Parameterization and preparation of lipidated LC3 on membrane

To investigate the dynamics of lipidated LC3 family proteins, we modeled LC3 conjugated with a phosphoethanolamine (PE) lipid anchor within the membrane. The methodology for attaching the PE anchor to the glycine residue follows previously published protocols [27]. For the coarse-grained (CG) model parametrization of the modified residue, glycine linked with PE (GLP), parameters were derived from the existing MARTINI force field for 1-palmitoyl-2-oleoyl-sn-phosphoethanolamine (POPE). The atomistic structure was converted into a CG model using the MARTINI force field and the martinize script, obtained from the MARTINI website . This script was modified to incorporate the parameters and bead definitions for the GLP residue. The protein structure was defined using standard MARTINI CG parameters, complemented by an ElNeDyn elastic network model to maintain structural integrity during simulations [49,50].

### Generation of heterogenous membrane model

A mixture of the most prevalent lipids found in endoplasmic reticulum (ER) was used to mimic lipid bilayer for depicting the autophagosome membrane [51]. The heterogeneous membrane used in this work is based on ER membrane composition i.e., DOPC 65%; DOPE 20%; POPI 10%; DOPS 5% with lipids distributed symmetrically between the inner and outer leaflets. The membrane was generated using the insane.py python script [52].

### Starting structure details

The initial structural system for studying the self-assembly of lipidated LC3 on a membrane comprised 90 lipidated LC3 proteins embedded within a 100 nm-wide heterogeneous membrane, designed to mimic the autophagosome environment. The PE anchor attached to the glycine residue of each protein was inserted into the phosphate headgroup region of the membrane to facilitate embedding. To maintain a biologically relevant orientation, the proteins were positioned such that their four previously identified membrane-interacting regions remained in close proximity to the membrane [26]. The proteins were initially spaced approximately 8 nm apart from one another, ensuring an even distribution while allowing for interactions during the simulation.

### Protocol for coarse grained molecular dynamics simulations

CG MD simulations were performed using the MARTINI model (version 2.2) with the MARTINI2.2 force field [49]. We first constructed initial CG model of the lipidated LC3 consisting of 291 beads for 120 residues. DSSP secondary structure assignments were used to generate backbone restraints that preserve local secondary structure for models [53]. Elastic network has been applied to maintain the structural fold of the protein while keeping their internal flexibility. A single CG model of lipidated LC3 was embedded into the bilayer spanning the periodic simulation box in the xy plane. The initial configuration and parameters of the CG model of lipidated LC3 was first tested by running a short simulation of 1 μs prior to simulating multiple copies. The resulting system consists of 90 lipidated LC3, with 26,190 beads and 33280 lipids. Each simulation was performed for 10 μs and a total of 30 μs of simulation time from three replica runs was generated.

All simulations were performed using Gromacs 2020 and the standard MARTINI protocol [54]. Periodic boundary conditions were employed, and three simulations were run with a time step of 20 fs, one with a time step of 10 fs producing 10 μs of data each. The system was solvated with the standard MARTINI water model using the insane.py script [55]. The system was first minimized and equilibrated using the Berendsen thermostat and barostat along with position restraints on protein backbone beads followed by production runs with a 20 fs time step [56]. System temperature and pressure during the production phase were maintained at 310 K and 1atm with the velocity rescaling thermostat and the semi-isotropic Parrinello–Rahman barostat, respectively [57]. The van der Waals interactions cutoff of 1.2 nm was used with a switching function applied from 0.9 nm in all simulations, and the reaction field coulomb type was utilised with a switching function from 0.0 to 1.2 nm. To constrain the covalent bonds LINCS algorithm was applied [58].

### Simulation analysis

We conducted clustering analysis using custom Python scripts using essential libraries such as NumPy, MDAnalysis. To identify oligomerization interfaces or sticky patches, we analyzed the coarse-grained interaction density of LC3 proteins. For multimeric LC3 simulations, we calculated the average occupancy, representing the percentage of time frames during which each interacting pair was observed across all simulations. In clustering algorithm, protein interactions were defined based on the proximity of their centers of geometry. Proteins were considered interacting if their centroids were within a 5 nm radius. Since the LC3 protein has an approximate diameter of 5 nm, molecules within this range were treated as potential interactors. Additionally, neighbouring proteins were determined to interact when the center of geometry of their residues were within 0.1 nm of each other. This dual metric allowed us to robustly capture interaction dynamics and cluster formation within the simulation environment. To analyze the oligomerized population of LC3 within the 90 chains, we first calculated the distances between the phosphate beads of the GLP residues across all possible chain combinations. Specifically, we examined pairwise distances among all 90 LC3 chains to identify potential interactions. Chains were flagged as interacting when the distance between their phosphate beads was below a critical threshold of 5 nm, which served as the basis for characterizing oligomerization.

To further identify and characterize interaction interfaces, residue occupancy contact maps were generated for all interacting pairs. These maps were constructed using the last 5 microsecond of the simulation trajectory, offering a high-resolution view of interaction dynamics. A contact between two residues was defined when the distance between them was within 8 Å and the occupancy exceeded 30% of the simulation time. This detailed approach allowed us to precisely map the spatial and temporal features of the LC3B oligomerization interfaces. We combined the contact map matrix from all the generated pairs to construct a cumulative contact map matrix. This cumulative matrix was then normalized by dividing it by the total number of contacts formed.

MDAnalysis tools were extensively utilized for various analyses, such as contact analyses, protein-membrane interactions, and lipid 2D density calculations [59]. The LiPyphilic package was employed to determine the enrichment index for each lipid at a cutoff of 0.12 nm [60]. Molecular visualizations were generated using PyMOL and ChimeraX.

### All-atomistic simulation protocol for LC3 dimer

The final snapshots from all three simulations, with the clustered population of LC3, were backmapped to all-atom resolution using the *backward.py* MARTINI script [61]. Each dimer interface was subsequently energy-minimized to resolve any steric clashes. Pocket volumes for each LC3 molecule in the three simulations were calculated using POVME 3.0 on the static backmapped structures [62]. The center of geometry was determined separately for the rear-side where Loop6 forms the pocket, as well as the HP1 and HP2 pockets, and used as the reference coordinates for volumetric calculations. A probe size of 0.6 nm and a grid spacing of 0.05 nm were employed to increase the accuracy of the volume measurements.

To investigate the impact of the Loop6-α2 region of the LC3 clustering interface on LC3 self-assembly, four mutants were designed and analyzed. The mutations were introduced in the top-ranked binding poses of the LC3 dimer interface, identified in the CG-MD simulations using PyMOL. The mutated protein structures were subsequently positioned on a membrane composed of heterogeneous lipids. Control simulations of WT LC3 in the same dimer orientation were also performed for comparison. Dimer simulations of wild-type (WT) and mutant LC3 structures were prepared using the CHARMM36m force field (2019 version) [63], while previously validated parameters were employed for GLP residue generating total simulation time of 8 μs.

### Cell Culture and Immunofluorescence

BS-C-1 cells, African green monkey (*Cercopithecus aethiops*) kidney epithelial cells (CCL-26; American Type Culture Collection) were maintained in a complete growth medium with 10% (v/v) fetal bovine serum (FBS), 2 mM L-glutamine, 1 mM sodium pyruvate, and penicillin-streptomycin. Cell culture condition was maintained at 37°C with 5% carbon dioxide. BS-C-1 cells were seeded on #1.5 coverslip bottom 35mm confocal dishes (Ibidi) or eight-well Lab-Tek chambers (Ibidi) for TIRF and STORM imaging. For inducing autophagy, cells were starved by incubating in Krebs-Ringer’s solution (pH 7.4) (KRBH), supplemented with 4.5 mM glucose, 0.1% BSA, and 1 mM sodium pyruvate for 2.5 hours at 37°C [64]. For immunofluorescence, cells were fixed with absolute ice-cold methanol for 7min at -20°C. Post fixation, cells were rinsed with PBS twice, 2 min each at room temperature, followed by blocking for 2 hours at room temperature in blocking buffer (3% w/v BSA with 0.2% Saponin in PHEM buffer (w/v; Sigma Aldrich). Cells were then incubated with primary antibodies diluted in blocking buffer overnight at 4°C, followed by two washes of 5 min each with 1X PBS (Gibco). Subsequently, cells were incubated with AF647-labeled secondary antibodies for 1 hour at room temperature, followed by three washes of 5 min each with 1X PBS.

Transient transfection of plasmids (Table S1) was performed using XtremeGene HP DNA transfection reagent (Roche). Briefly, the plasmid DNA and transfection reagent were mixed in serum-free media and kept at room temperature for 20 mins. The final volume was adjusted with complete MEM, and the transfection was carried out for 24 hrs.

### STORM Imaging Protocol

2D Super-resolution STORM images were acquired with a custom-built microscope. A 647 nm continuous wave visible fiber laser (MPB communications) was used to excite Alexa Fluor 647 (Invitrogen), while a 405-nm solid-state laser (Obis; Coherent) was used for reactivating the Alexa Fluor 647 (Invitrogen) via an activator dye (Alexa Fluor 405). The emitted light from Alexa Fluor 647 (Invitrogen) was collected by the Nikon 100× oil immersion TIRF-SR objective (NA 1.49), filtered by an emission filter (ET705/72m; Chroma), and imaged onto the EM-CCD camera (20 ms per frame), with a total of ∼60,000 frames acquired per image. For multicolour STORM imaging, we performed sequential imaging with AF647 dye. We immunostained for LC3 and performed STORM imaging as described above, followed by treatment with sodium borohydride (1mg/mL) treatment for 7 min at room temperature to permanently quench the AF647 signal from LC3. Next, we performed immunostaining in situ on the microscopic stage for PI3P via mCherry antibody using AF647. STORM images of both channels were aligned by matching the localization obtained from the fiduciary markers (TetraSpeck microspheres, Invitrogen T7279)., using a custom image registration software, as reported before [65]. 3D dSTORM imaging was done with a SAFeRedSTORM module (Abbelight) mounted on an Evident/Olympus IX3 microscope having an oil-immersion 100x objective (1.5NA oil immersion, Evident/Olympus) and fiber-coupled 642nm laser (450mW Errol). Fluorescent signal of single molecules was collected with an ORCA-Fusion sCMOS camera (Hamamatsu). A cylindrical lens was placed before the camera to introduce astigmatism. Image acquisition and control of microscope were driven by Abbelight’s NEO software. Each image stack contained 60,000 frames. Selected ROI (region of interest) dimension was 512 × 512 pixels (pixel size 97 nm). Cross-correlation was used to correct for lateral drifts. Super-resolution images (.tiff) with a pixel size of 10 nm and localization files (.csv) were obtained using NEO_analysis software (Abbelight).

### Cluster analysis

Segmentation of localizations in STORM images was performed using the density-based clustering algorithm DBSCAN [66]. DBSCAN requires two parameters: the minimum number of points (k) within a search radius (epsilon). A threshold of a minimum of 12 localizations was applied to remove the noise. To determine the epsilon in an unbiased manner, the elbow plot method was used. For cluster analysis, ROI (region of interest) containing single autophagosomes was cropped. To estimate the size of LC3 clusters identified through DBSCAN analysis, the centroid of each cluster was calculated by taking the geometric mean of all localizations within the cluster. The outer boundary of each cluster was then approximated using a convex hull, which connects the farthest localizations within the cluster [67]. Subsequently, the distance from the centroid to the convex hull boundary points was measured for each cluster, and the mean of these distances was taken as the cluster radius. This method provides a robust estimation of cluster size by incorporating both the centroid and edge boundaries defined by the spatial distribution of localizations. For comparative analysis of the LC3 clustering between WT and mutants, we enumerated LC3 area occupancy on autophagosome (Cluster area: Total area) and number of LC3 nanoclusters per autophagosome.

### Single-Particle Tracking

To perform single-particle tracking analysis of autophagosomes, cells were transfected with LC3 plasmid constructs for both WT and mutant variants. Following transfection, cells were starved as described previously and subsequently subjected to live-cell imaging using TIRF microscopy. Imaging was performed at a 100 ms time interval for 100 sec. Raw data for autophagosome motility were processed by background subtraction using a rolling ball radius of 50.0, followed by bleach correction through histogram matching. The processed data were then analyzed using the TrackMate plugin [68] in Fiji software with the parameters listed in Table S3. Data were exported to Microsoft Excel (2013) and Origin 2022b for further analysis.

### Statistical analysis

For the quantification of microscopy experiments, the statistical significance of differences between WT and mutant samples was assessed using the non-parametric Mann–Whitney U test. Significant differences were denoted as * for p ≤ 0.05, ** for p ≤ 0.01, and “ns” for non-significant results.

## Acknowledgments

LT is thankful to the funding support from the DBT/Wellcome Trust India Alliance (IA/21/2/505925). NM acknowledges funding support from the DST-SERB (CRG/2023/008391) and Indian Council of Medical Research (ICMR – 2021-15239/GTGE/Adhoc-BMS). SF was supported by ICMR-SRF fellowship. DMK was supported by the Prime Ministers Research Fellowship (PMRF). The authors thank all members of the lab for their support. We thank Prof. Melike Lakadamyali (University of Pennsylvania, Philadelphia, PA) and Prof Bo Huang (University of California, San Francisco) for the STORM analysis software. We are also thankful to Dr. Benjamin Compans and Abbelight for helping with 3D STORM imaging and analysis software.

