## Supplementary File for "LC3 forms functional nanocluster on autophagosome"

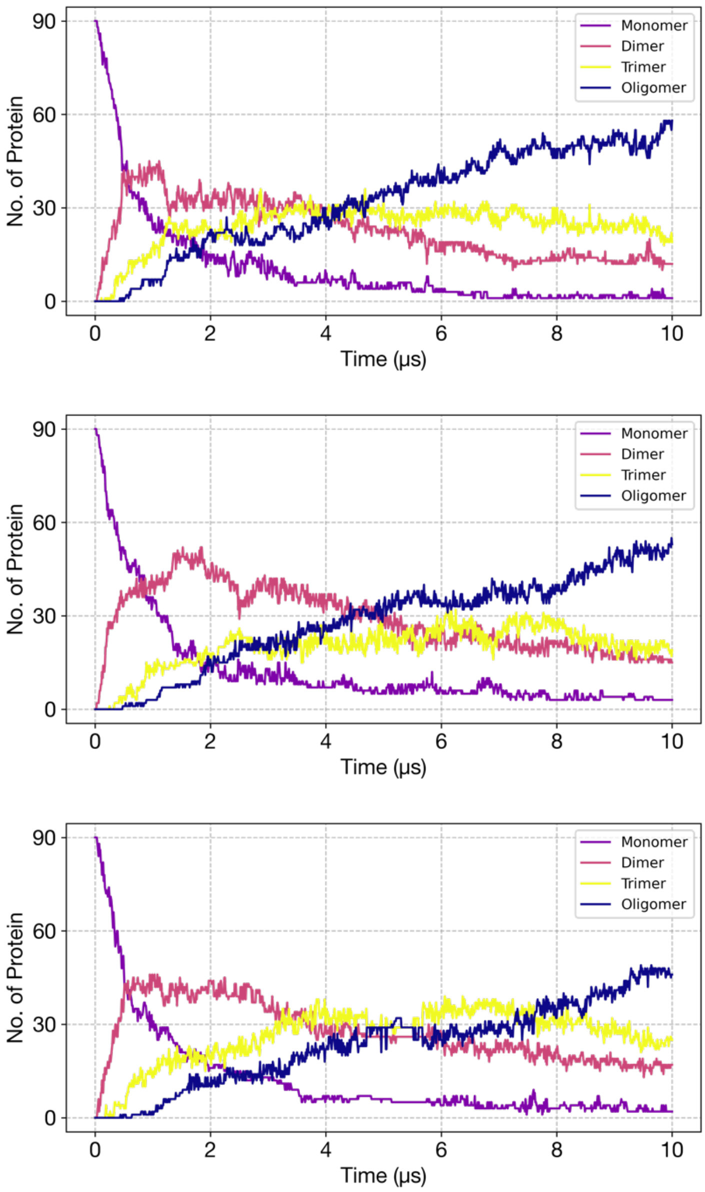


**Figure S1. Time evolution of oligomer formation across three trajectories.** The coarse-grain simulations started with 90 proteins arranged in a grid-like configuration. Over the course of the simulations, dimers and trimers initially formed, followed by the emergence of higher-order LC3 oligomers within 3 μs.

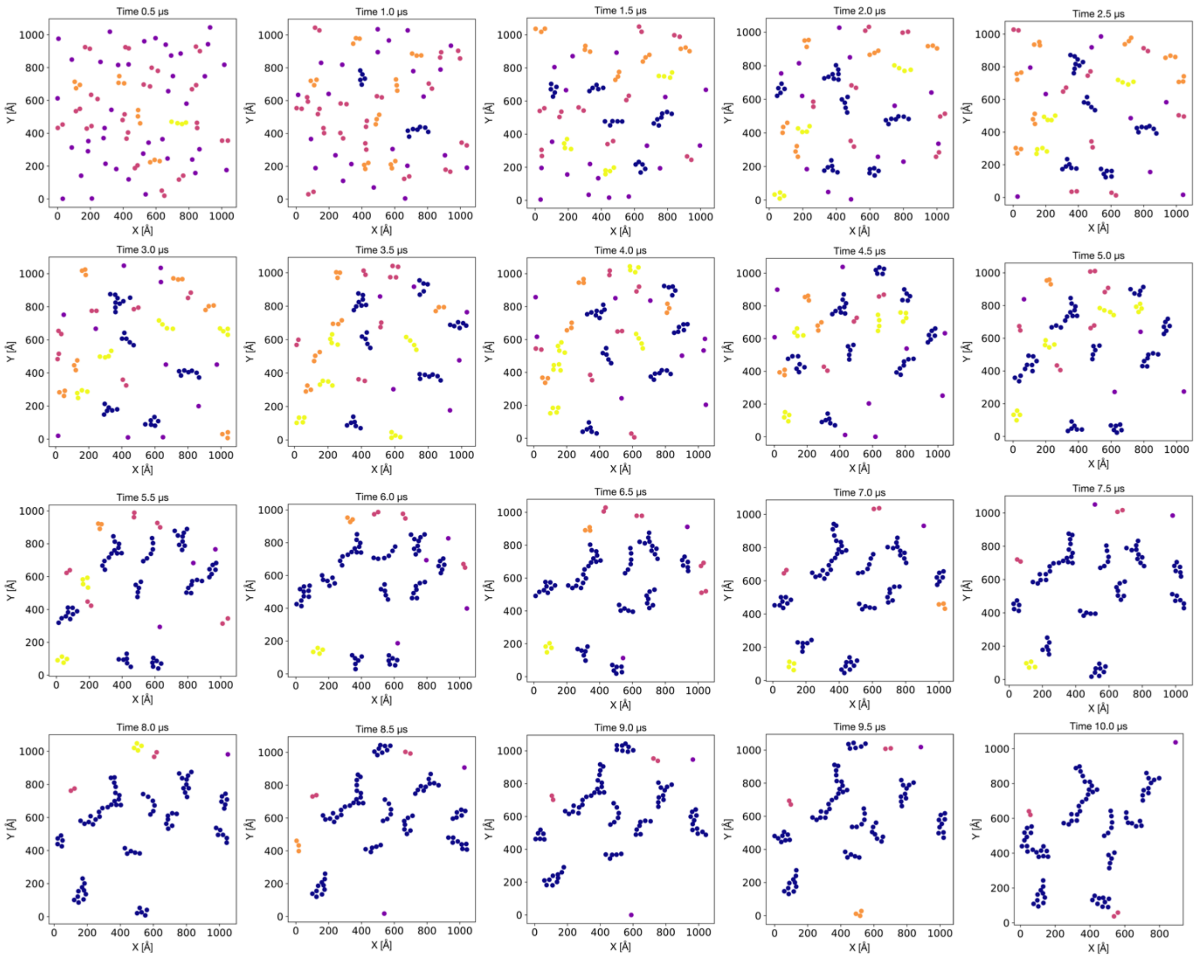


**Figure S2. Visualization of LC3 cluster formation over time across 10 μs trajectory.** The positions of LC3 molecules are plotted relative to the dimensions of the simulation box at 500 ns intervals to track the progression of cluster formation. Circle colors represent cluster types: purple (monomer), pink (dimer), orange (trimer), yellow (tetramer), and dark blue (oligomer). The cluster distribution is displayed for all three simulation replicates.

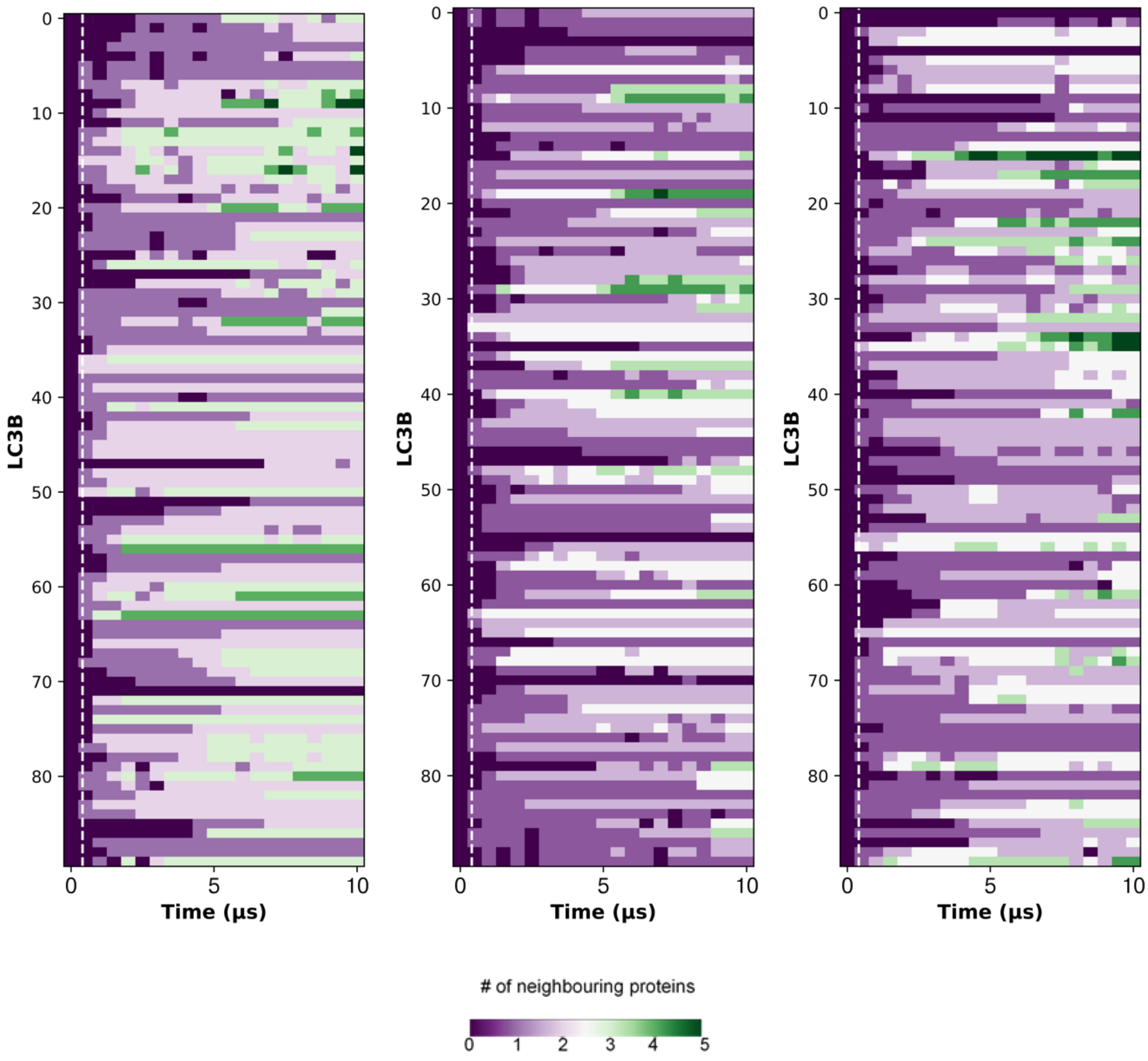


**Figure S3. The heatmap shows neighbouring proteins in LC3B clusters.** Each of the three boxes represents a separate simulation, with each row within a box corresponding to an individual protein over the simulation time. The color scale indicates the number of neighbouring proteins, defined by a 5 nm cutoff distance between the centers of mass of two proteins. Neighbor counts range from 0 (purple) to 6 (green).

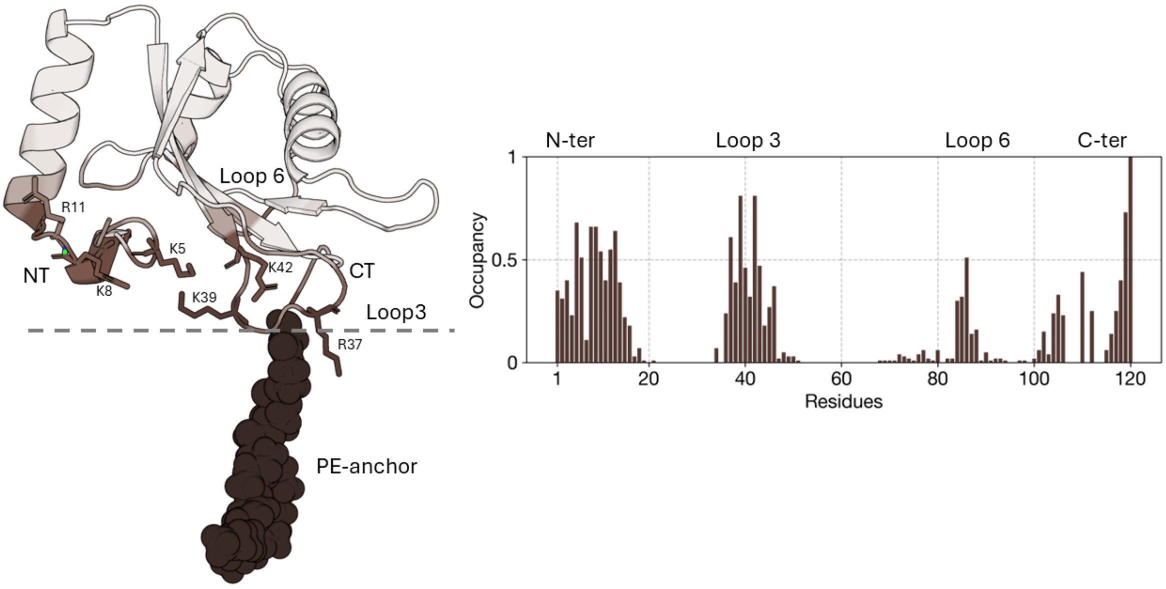


**Figure S4. Membrane-interacting module (MTM) in the clustered population of LC3.** Residues interacting with the membrane are mapped onto a representative structure, with color intensity reflecting interaction occupancy during the final 5 µs of the simulation trajectory (interaction cutoff distance was 5 Å). The accompanying bar plot shows the average occupancy values for each residue across all 90 chains.

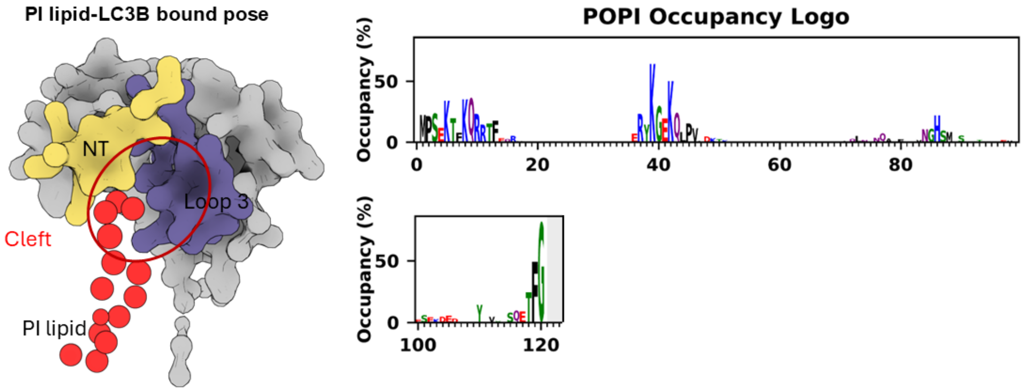


**Figure S5. Representative bound pose of a phosphatidylinositol (PI) lipid species interacting with the LC3B protein.** The selected top-ranked pose, identified through back-mapping from coarse-grained simulations, reveals binding within a cleft formed between Loop 3 and the N-terminal region of LC3B. The accompanying logo plot depicts the amino acid composition of the binding site, which is predominantly enriched with positively charged residues.

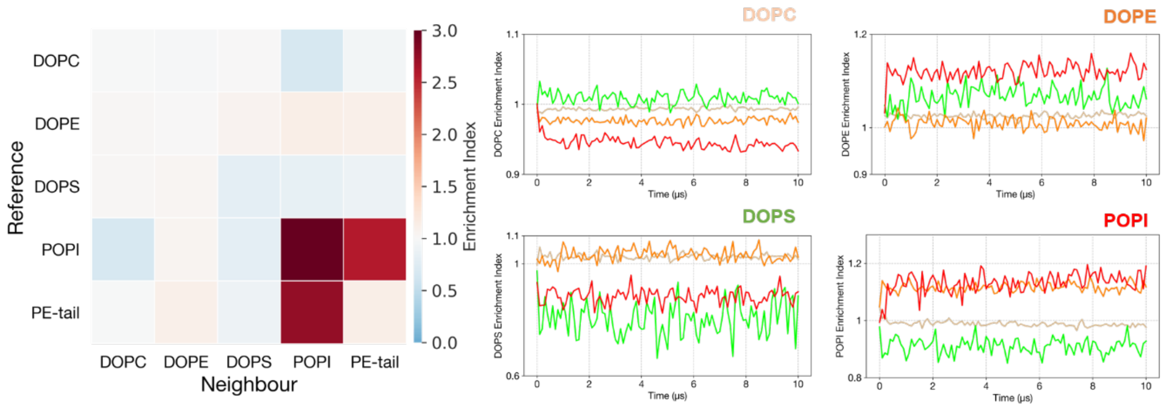


**Figure S6. Lipid enrichment around LC3 clusters.** Heatmap showing lipid presence and depletion around LC3B, with the enrichment and depletion index for each lipid species calculated by analyzing the number of lipids within a 12 Å distance from LC3B in each frame. Index values above 1 indicate lipid enrichment, while values below 1 indicate lipid depletion. The heatmap emphasizes the dynamic clustering and distribution of POPI lipid species near LC3B. The accompanying line plot depicts the self-enrichment index of POPI lipids over time, providing a temporal perspective on lipid dynamics.

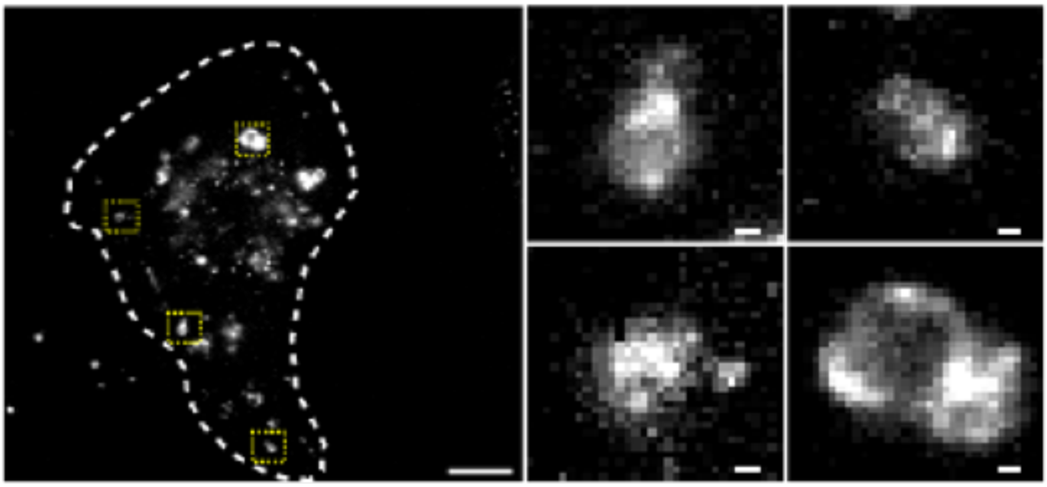

**Figure S7. TIRF imaging of autophagosome marked by LC3 immunostaining**: Conventional TIRF microscopy image of autophagosomes reveals LC3 localised to autophagosome recapitulate autophagosome structure, traditionally referred as autophagosome puncta. The diffraction limited TIRF imaging does not resolve the individual LC3B proteins localised on autophagosome. Scale bar 10 µm (image), 500 nm (zoom).

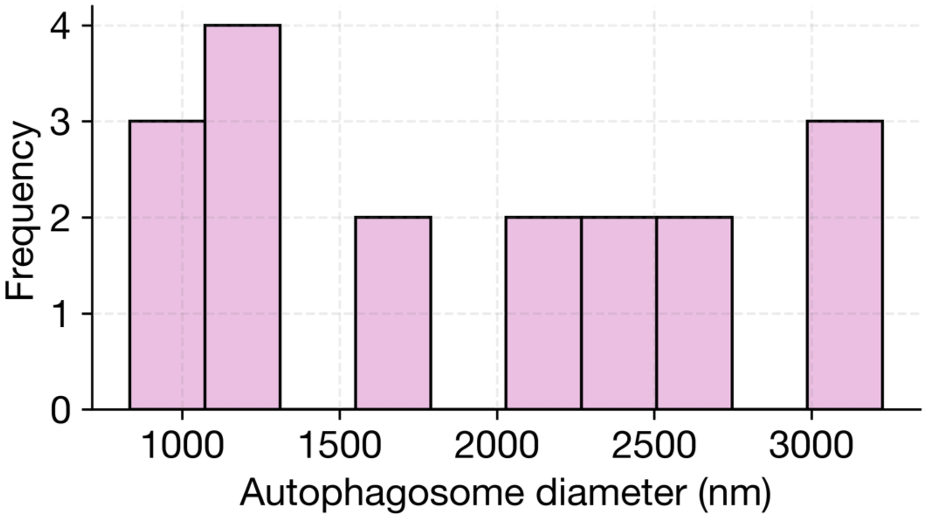


**Figure S8. Quantification of autophagosome size from STORM Imaging**. Histogram showing the distribution of autophagosome sizes, highlighting their variability.

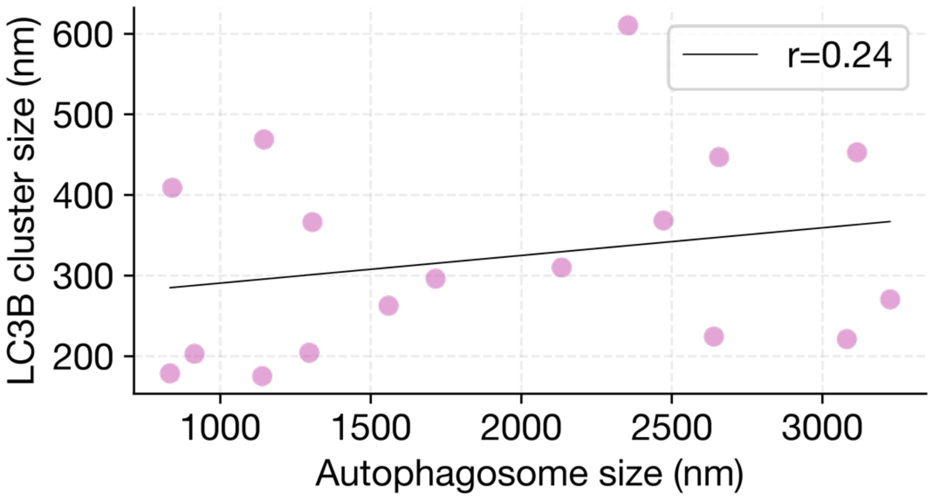


**Figure S9. Correlation analysis between autophagosome size and LC3B cluster size, revealing a non-linear relationship.** The average size of LC3 clusters does not proportionally vary with autophagosome size variability.

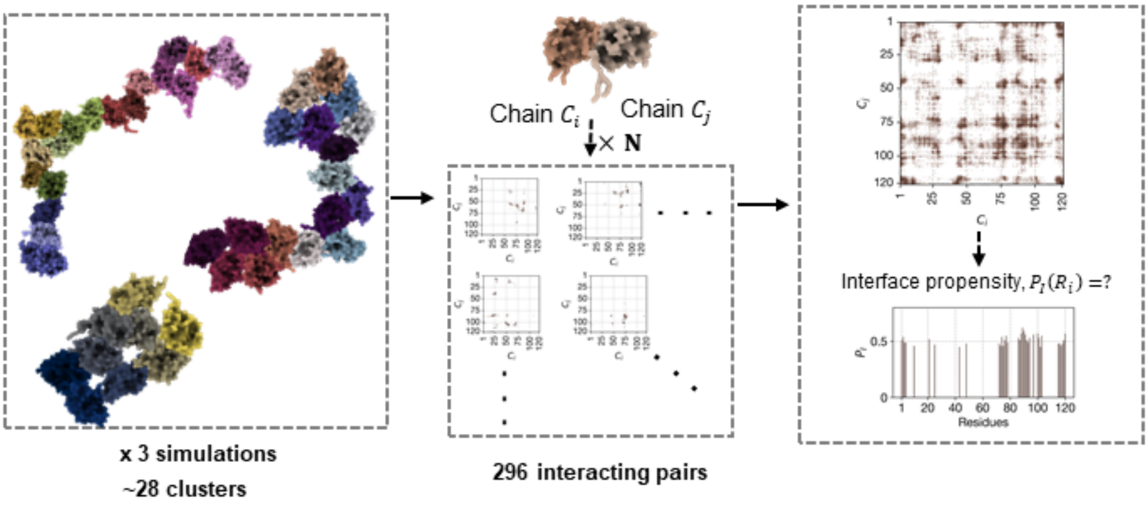


**Figure S10: Schematic workflow illustrating the process of identifying clustering interfaces.** From 28 LC3B homo-oligomers obtained through MD simulations, encompassing 296 dimer interfaces. To analyze region-specific interaction propensities, contact maps were generated for each residue within each cluster, highlighting the likelihood of individual residues participating in the clustering interface.

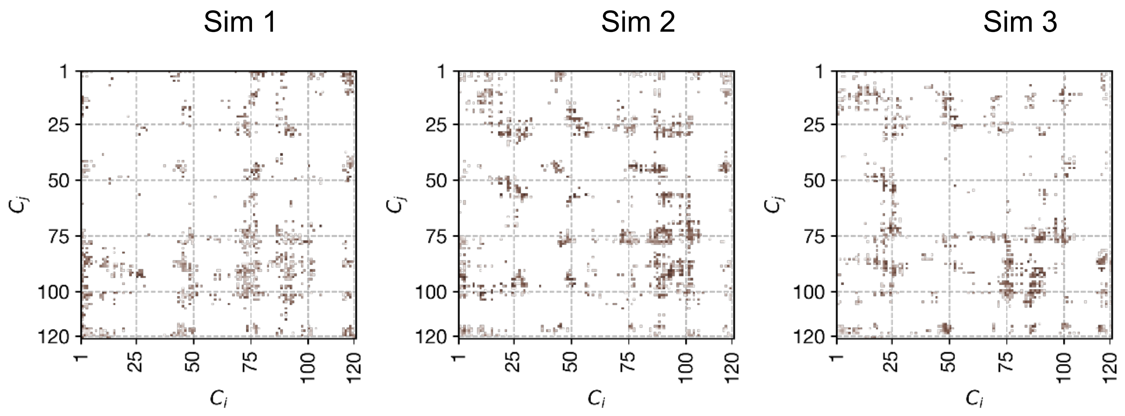


**Figure S11. Contact matrices for all residue pairs across three simulations.** Cumulative contact map showing contacts with an occupancy greater than 0.3 across all interfaces observed in each simulation. The color gradient represents occupancy, ranging from 0 to 1.

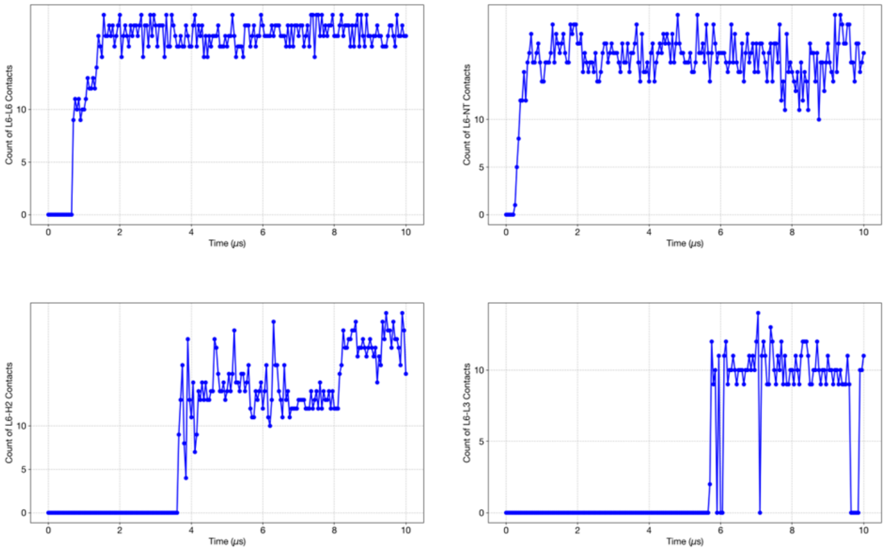


**Figure S12: Time evolution of the top binding poses interactions forming the interface.** The most probable region to be at the interface i.e, Loop 6 formed interactions early on the simulation leading to the clustering. Other regions which constitute the periphery of the interface formed interactions later.

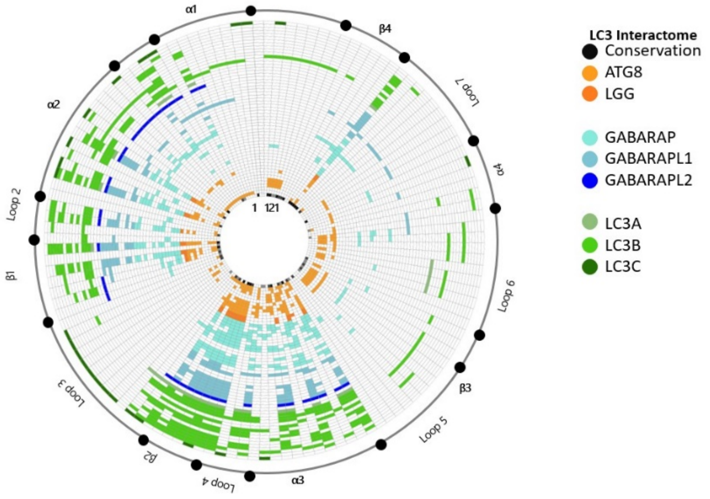


**Figure S13: LC3 protein-protein interaction map at residue level derived from experimental complexes.** A total of 46 bound crystal structures of all the orthologs of LC3 protein were included for understanding interfacial regions of LC3 complexes. The interface residues were calculated by taking a cutoff of 5Å. Radial plot displaying residue-wise information from available bound crystal structure data of all the orthologs of LC3B, illustrating the participation of each residue at the binding site.

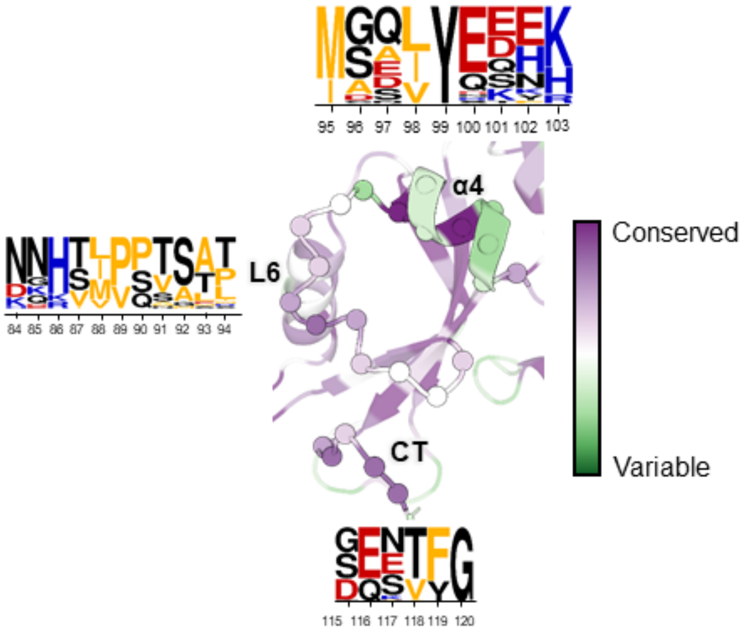


**Figure S14. Conserveness of rear binding pocket.** The representative snapshot highlights the novel rear-side Loop6 pocket formed by Loop6-α4 in LC3B, identified as a key clustering interface alongside CT residues. Conservation scores for this region are mapped onto the structure, providing insights into evolutionary conservation. A corresponding logo plot illustrates the residue distribution at each site, emphasizing the conserved nature of this pocket.

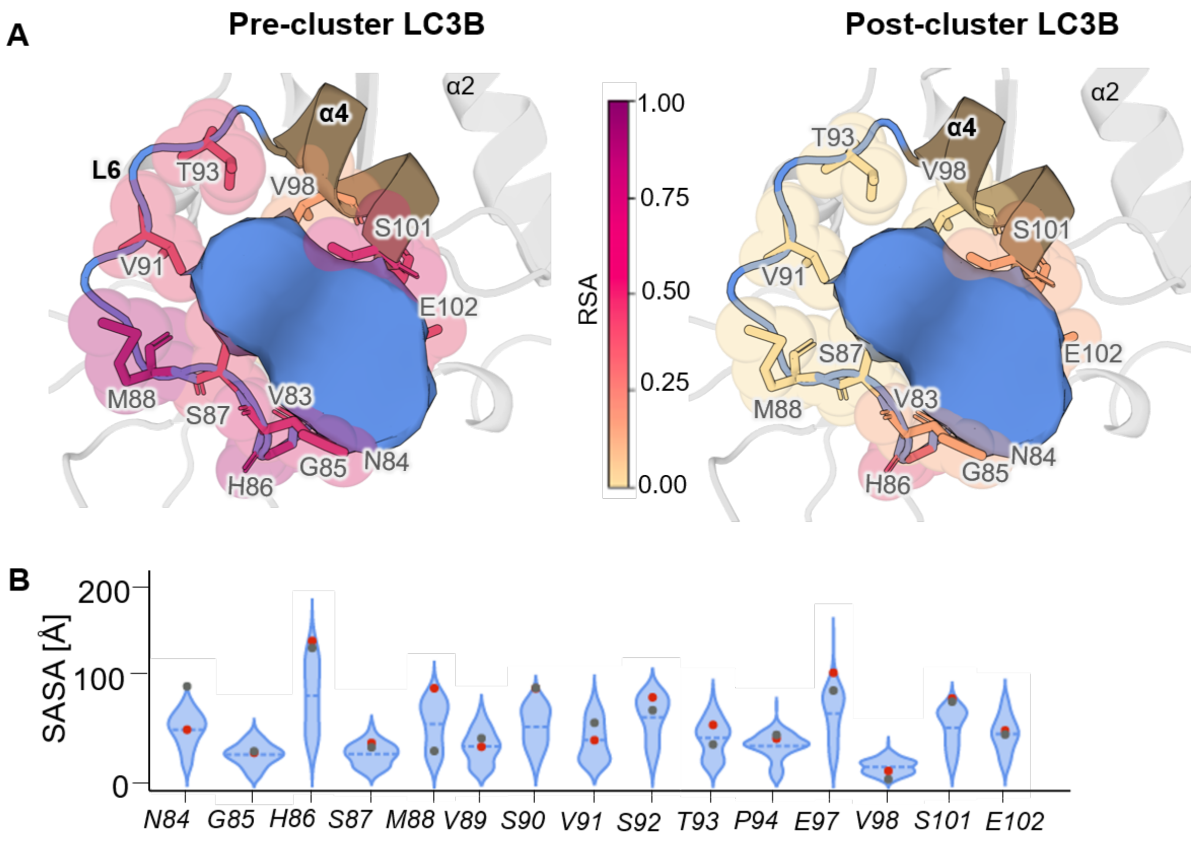


**Figure S15. Snapshots highlighting the relative residue accessibility in the most probable interface region between Loop 6 and α4 of LC3B.** A) Residue-specific relative solvent accessibility (RSA) differences (ΔRSA, unbound - bound) < 25% are mapped to emphasize their contribution to binding. (B) Violin plot showing the distribution of solvent-accessible surface area (SASA) for interface residues in LC3B dimers, demonstrating that most Loop 6 residues remain fully or partially exposed to facilitate clustering.

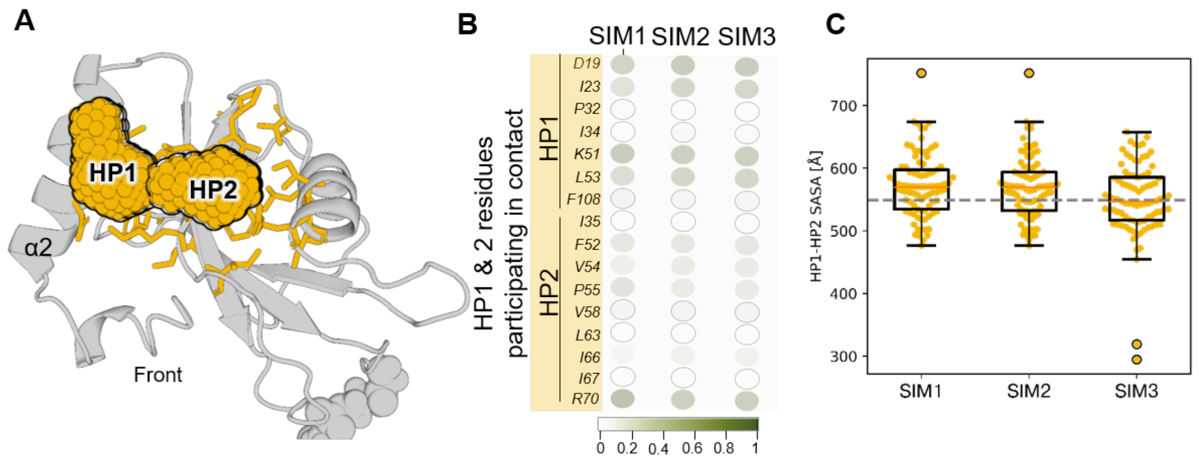


**Figure S16. Canonical binding pocket is non-overlapping with LC3B clustering interface.** (A) The two canonical hydrophobic pockets, HP1 and HP2, are highlighted in mustard color. (B) Binding pocket residues were analyzed for interactions contributing to clustering, but no such contacts were observed. (C) Surface accessibility calculations confirmed that the binding pocket remains exposed and available for potential binding.

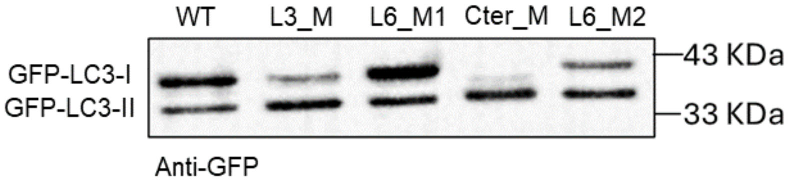


**Figure S17. Presence of two populations of LC3.** Western blot analysis of GFP tagged LC3B wildtype protein and four mutant proteins was performed to confirm the lipidation.

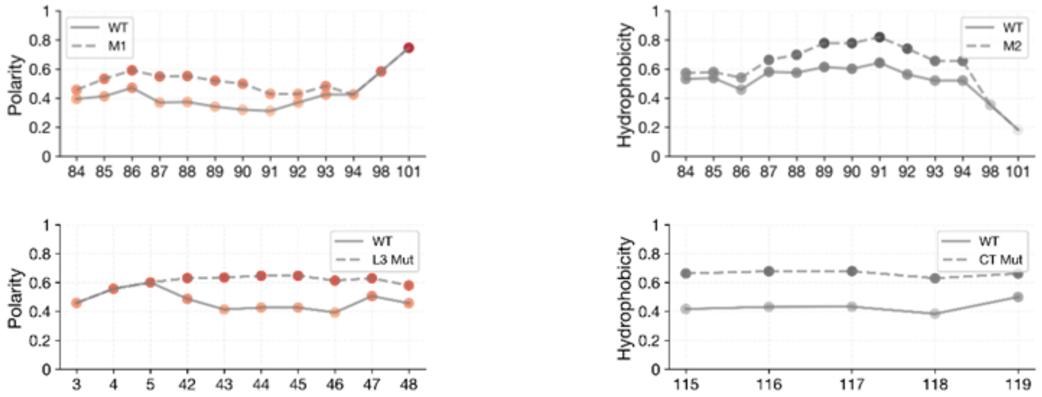


**Figure S18.** Quantitative plot shows the differences in polarity and hydrophobicity based on the Grantham and Kyte & Doolittle scales for each mutant.

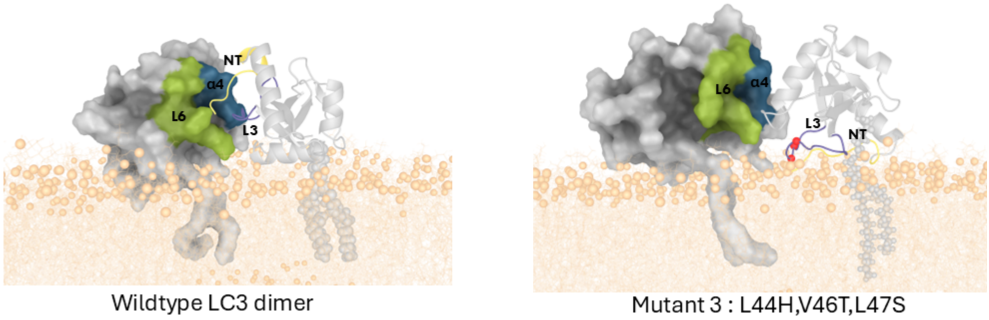


Wildtype LC3 dimer

Mutant 3 : L44H,V46T,L47S

**Figure S19.** Final configurations of Loop3 mutants after 650 ns of molecular dynamics simulations, alongside the wild-type (WT) in the same starting configuration. The Loop 3 mutant exhibited no dissociation between LC3B molecules compared to the Loop 6 mutants but a change in the orientation of the interface residues was observed.

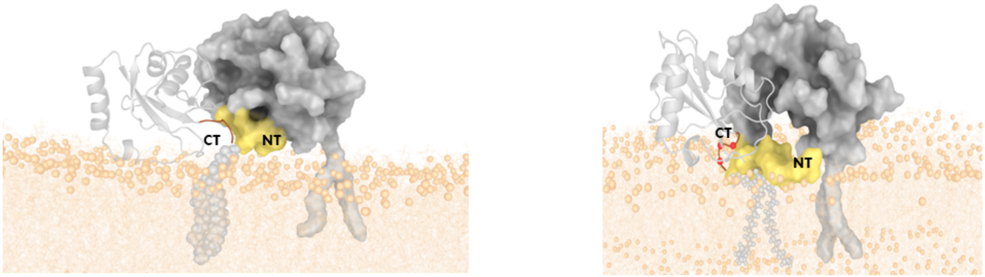


Wildtype LC3 dimer

Mutant 4 : Q116A,E117A,T118A

**Figure S20.** Final configurations of C-terminal mutants after 650 ns of molecular dynamics simulations, alongside the wild-type (WT) in the same starting configuration. The C-ter mutant exhibited no dissociation between LC3B molecules.

**Table S1: Primary and Secondary antibodies used in study.**

| Sr. No. | Antibody | Identifier | Dilution used |
|---|---|---|---|
| 1 | Rabbit Anti-LC3B | CST #3868 | 1:100 |
| 2 | Chicken Anti-mCherry | NBP2-25158 | 1:500 |
| 3 | ChromoTek GFP-Booster Alexa Fluor® 647 | gb2AF647 | 1:1000 |
| 4 | Anti-rabbit IgG-AF647 | 711-005-152; Jackson ImmunoResearch | 1:100 |
| 5 | Anti-Chicken IgG-AF647 | 703-005-155; Jackson ImmunoResearch | 1:100 |

**Table S2: Plasmids Used in this study.**

| Sr. No. | Plasmid ID | Identifier |
|---|---|---|
| 1 | pcDNA3.1-GFP-LC3B-WT | Custom made |
| 2 | pcDNA3.1-GFP-LC3B-L3_M | Custom made |
| 3 | pcDNA3.1-GFP-LC3B-Cter_M | Custom made |
| 4 | pcDNA3.1-GFP-LC3B-L6_M1 | Custom made |
| 5 | pcDNA3.1-GFP-LC3B-L6_M2 | Custom made |
| 6 | pmCherry-2xFYVE | Addgene#140050 |

**Table S3: Parameters used for Single Particle Tracking (SPT) analysis in TrackMate plugin**

| Parameters | Value |
|---|---|
| Vesicle diameter | 0.6 µm |
| Detector | LoG (Laplacian of Gaussian) |
| Initial thresholding | None |
| Tracker | Simple LAP tracker |
| Linking max distance | 0.3 µm |
| Gap-closing max distance | 1 µm |
| Gap-closing max frame gap | 20 |
| Filter on track | Number of spots in track |
